# Loss-of-function cancer-associated mutations in the EIF4G2 non-canonical translation initiation factor

**DOI:** 10.1101/2023.08.22.554280

**Authors:** Sara Meril, Marcela Bahlsen, Miriam Eisenstein, Alon Savidor, Yishai Levin, Shani Bialik, Shmuel Pietrokovski, Adi Kimchi

## Abstract

Tumor cells often exploit the protein translation machinery, resulting in enhanced protein expression essential for tumor growth. Since canonical translation initiation is often suppressed due to cell stress in the tumor microenvironment, non-canonical translation initiation mechanisms become particularly important for shaping the tumor proteome. EIF4G2 is a non-canonical translation initiation factor that mediates internal ribosome entry site [IRES] and upstream open reading frame [uORF] dependent initiation mechanisms, which can be used to modulate protein expression in cancer. Here we explored the contribution of EIF4G2 to cancer by screening the COSMIC database for EIF4G2 somatic mutations in cancer patients. Functional examination of missense mutations revealed deleterious effects on EIF4G2 protein-protein interactions, and importantly, on its ability to mediate non-canonical translation initiation. Specifically, one mutation, R178Q, led to reductions in protein expression and near complete loss-of-function. Two other mutations within the MIF4G domain specifically affected EIF4G2’s ability to mediate IRES-dependent translation initiation but not that of target mRNAs with uORFs. These results shed light on both the structure-function of EIF4G2 and its potential tumor suppressor effects.

## Introduction

Eukaryotic mRNA translation is a highly regulated multistep process that regulates protein synthesis under normal and stress conditions. mRNA translation is largely divided into three steps-initiation, elongation and termination (Jackson *et al*, 2010). In canonical cap-dependent translation, the initiation step involves the recruitment of eukaryotic initiation factors of the multi-protein complex EIF4F on the mRNA 5’ cap m^7^GTP structure [m^7^G cap]. EIF4F is comprised of the cap binding EIF4E, the helicase EIF4A, and EIF4G1, which acts as a scaffold bridging the aforementioned factors, as well as PABP and EIF3, to enable 40S ribosome recruitment (Jackson *et al*., 2010). Notably, non-canonical mechanisms of translation initiation have also been described. One of the main players in these non-canonical mechanisms is EIF4G2 (also known as DAP5/p97/Nat1), a member of the EIF4G family of initiation factors (Imataka *et al*, 1997; Levy-Strumpf *et al*, 1997; Yamanaka *et al*, 1997). Unlike EIF4G1, EIF4G2 lacks the PABP and EIF4E binding sites, and therefore instead participates in several cap-independent modes of translation initiation, such as those utilizing internal ribosome entry sites [IRES] (Henis-Korenblit *et al*, 2002; Hundsdoerfer *et al*, 2005; Lewis *et al*, 2008; Liberman *et al*, 2015; Marash & Kimchi, 2005; Warnakulasuriyarachchi *et al*, 2004) or cap-independent translation enhancers [CITE] (Haizel *et al*, 2020), and N6-methyladenosine [m6A] driven translation (Yang *et al*, 2017). Additionally, EIF4G2 has been shown to promote read-through of 5’ upstream open reading frames [uORFs] (Smirnova *et al*, 2022) and/or re-initiation of the main coding sequence [CDS] following cap-dependent translation of these uORFs (Weber *et al*, 2022). It can also mediate a non-canonical cap-dependent method of initiation by binding EIF3D (de la Parra *et al*, 2018a; Volta *et al*, 2021), which replaces EIF4E as the mRNA 5’ cap interactor (Lee *et al*, 2016).

In order to maintain enhanced proliferation and altered cellular metabolism, tumors need to upregulate their protein translation capacity, and as such, cap-dependent translation becomes a convergent point for regulation in cancer cells (Smith *et al*, 2021). In fact, the involvement of the canonical initiation factors in cancer formation and progression is well established (de la Parra *et al*, 2018b). For example, EIF4E is often highly expressed in different cancers and can drive the translation of specific mRNAs such as VEGFA, MYC and TGFB1, which ultimately promote angiogenesis, cell proliferation and oncogenesis. Similar functions and cancer-promoting outcomes have been shown for overexpression of additional translation initiation factors such as EIF4A and EIF4G1 (de la Parra *et al*., 2018b). Yet, the tumor microenvironment is often characterized by cell stress, during which canonical cap-dependent translation is suppressed. In these circumstances, non-canonical translation initiation mechanisms, such as those involving IRESes and uORFs are utilized (de la Parra *et al*., 2018b; Smith *et al*., 2021). Yet while these specific mechanisms have been shown to be important for specific oncogene expression, such as MYC, little is known about how non-canonical translation initiation factors driving these mechanisms may contribute to cancer development and progression.

EIF4G2 is an appealing candidate to be studied in the context of cancer due to its established physiological roles. EIF4G2 is critical for embryonic development and differentiation of embryonic stem cells by driving non-canonical selective translation of critical mRNA cohorts (David *et al*, 2022; Yamanaka *et al*, 2000; Yoffe *et al*, 2016). Additionally, EIF4G2’s established translation targets during cell stress and apoptosis, including pro-apoptotic genes APAF1 and MYC, and anti-apoptotic IAP proteins (Henis-Korenblit *et al*., 2002; Liberman *et al*, 2008; Nevins *et al*, 2003; Warnakulasuriyarachchi *et al*., 2004), and BCL2, BCL2L1 and CDK1 during mitosis (Liberman *et al*, 2009; Marash *et al*, 2008), indicate its involvement in cell death and survival pathways. It is also involved in cellular responses to hypoxia and stress in various cancer cells, by mediating translation of PHD2 (Bryant *et al*, 2018) and an NH2-terminal truncated TP53 isoform from an internal IRES (Weingarten-Gabbay *et al*, 2014), respectively. These specific functions suggest that EIF4G2 may be critical for cell fate decisions in cancer, although its multiple and sometimes opposing functions make it difficult to predict whether EIF4G2 can promote or suppress tumor development and growth. To date only a small number of studies have examined a direct role for EIF4G2 in cancer. High EIF4G2 mRNA expression was associated with gastric cancer and metastatic triple negative breast cancer, correlating with decreased overall and metastasis-free survival (de la Parra *et al*., 2018a; Fu *et al*, 2022). In metastatic breast cancer cells, knock-down of EIF4G2 resulted in decreased cell migration in cellular invasion and wound healing assays, increased apoptosis upon loss of cell adherence. Injection of the KD cells into mice produced tumors with similar growth as control cells, but with decreased metastasis, invasiveness and angiogenesis (Alard *et al*, 2023). Consistent with these functional data, many of the EIF4G2 target mRNAs in metastatic breast cancer cells were associated with cell migration, invasion, the epithelial-to-mesenchymal transition and survival, such as integrins, vimentin, SNAIL1/2 and ZEB1 (Alard *et al*., 2023). It is thus likely that in breast cancer, EIF4G2 promotes metastasis through its ability to enhance migration, invasion and cell survival. On the other hand, EIF4G2 expression was observed to be reduced in bladder cancer, correlating with tumor de-differentiation and invasiveness (Buim *et al*, 2005). Thus, the limited data to date suggest that EIF4G2 may act as either an oncogene or a tumor suppressor, depending on the tumor context, and call for a broader and deeper investigation into whether EIF4G2 gain- or loss-of-function is indeed associated with cancer development and/or progression.

Here we screened the COSMIC database for EIF4G2 somatic mutations in cancer patients. We focused on the possible effects of single missense mutations on EIF4G2 protein structure and function, specifically on protein-protein interactions and translation initiation functions. Through analysis of these mutations, we have established the occurrence of loss-of-function mutations in EIF4G2 and separated the IRES-dependent initiation functions from uORF-dependent initiation functions, opening the door to understanding the phenotypic outcome of its loss-of-function on cancer progression and aggressiveness.

## Results and Discussion

### Screening for EIF4G2 mRNA expression and mutational burden in cancer patients

The TCGA database was screened in an unbiased manner for EIF4G2 mRNA expression in healthy subjects compared to patients harboring primary tumors (Fig EV1). The analysis was performed on 24 different cancer histology subtypes for which healthy subjects’ data was also available, taking into consideration that the healthy and primary tumor specimens were not necessarily sequenced from the same individual. Notably, nearly one third of the analyzed subtypes showed significant reduction in EIF4G2 mRNA expression (*i.e*., uterine corpus endometrial carcinoma [UCEC], bladder urothelial carcinoma [BLCA], kidney renal clear cell carcinoma [KIRC], prostate adenocarcinoma [PRAD], head and neck squamous cell carcinoma [HNSC], kidney renal papillary cell carcinoma [KIRP] and thyroid carcinoma [THCA]; 7/24 analyzed tumors), while two tumors (glioblastoma multiforme [GBM] and cholangio carcinoma [CHOL]) demonstrated a significant increase in EIF4G2 mRNA expression. Overall, these results demonstrate that EIF4G2 mRNA expression differs according to the tumor type.

In light of the inconclusive nature of the expression data, and considering that EIF4G2 is a long-lived protein under specific translation control (Kearse *et al*, 2019; Takahashi *et al*, 2005; Tang *et al*, 2017), whereby its mRNA expression is particularly limited in its ability to predict EIF4G2 protein expression, we sought to determine a more definitive indicator of EIF4G2 function in patient tumors. In the absence of publicly available informative, abundant data on protein expression, we instead examined somatic mutations derived from cancer patients to determine if they serve as gain-of-function or loss-of-function mutations, or affect the protein levels of EIF4G2.

Sequencing data of 411 EIF4G2 mutations from 369 different patients, all confirmed to be independent somatic mutations, were screened and collected from the COSMIC database. Of these 369 samples, 290 (79%) were found in carcinomas, 42 (11%) in melanomas, with the remainder identified in cancers of different histology types at low percentages (Fig 1A). Focusing on carcinoma, the most abundant histology type, 101 mutations were found in adenocarcinomas (∼35%), 32 (11%) in squamous cell carcinomas, and 21 (7%) in endometrioid carcinomas (Fig 1A). 248 of the mutations were situated in the coding sequence, 77% (191/248) of which were categorized as missense mutations (Fig 1B). Using data analyses and statistical methods that carefully identify primary unique mutations and calculate their significance using the expected probabilities of all possible mutations (Nuta *et al*, 2019), six hotspots (26/191, 13.6%) with a statistically significant occurrence were identified (Fig 1C). In addition, 10.5% (26/248) of the mutations in the EIF4G2 coding sequence predicted a complete loss-of-function due to out of frame deletion, insertion, or early stop codon (non-sense) mutations (Fig 1B). The distribution across cancer types of the predicted deleterious mutations and the significant missense mutations is shown in Fig 1D, and represents the number of patients with each annotated mutation according to the primary tumor site. This in-depth, accurate statistical analysis of numerous naturally occurring tumor primary sites revealed that EIF4G2 mutations are mostly found in cancers of the large intestine and endometrium, representing 22% and 14%, respectively, out of the deleterious and significant missense mutations presented. These results emphasize the possible biological importance of EIF4G2 in colon and endometrial tumors. Moreover, the complete loss-of-function mutations are an important indication that EIF4G2 protein is probably reduced in both quantity and function in some cancer patients.

**Figure 1.**
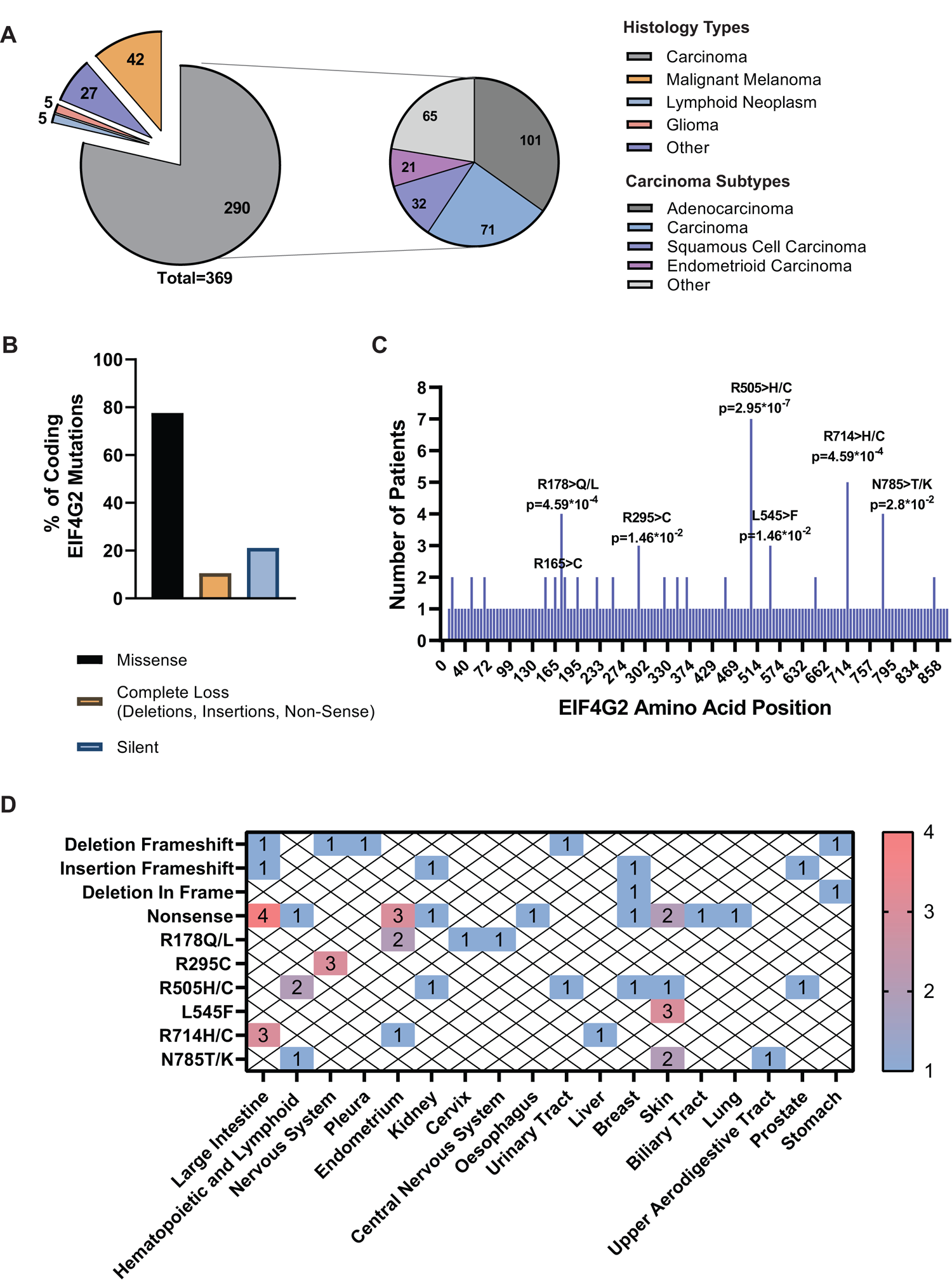
Distribution of mutations in EIF4G2 gene in cancer patients. **A** Pie chart representing the distribution of 369 patient samples with EIF4G2 mutations according to tumor histology type (Left). Pie-of-pie chart representing the tumor histology sub-type of the 290 carcinoma patients with EIF4G2 mutations (Right). **B** Bar graph showing the mutation classification in 248 EIF4G2 coding mutations. **C** Distribution and occurrence of 191 verified somatic missense mutations in the EIF4G2 coding region, from 248 independent tumor samples, as identified in the COSMIC database. Positions with significant mutation occurrences are labeled with the amino-acid substitution and *p*-value, calculated for windows of 1, 3, 9, 15, 30, & 60 bases, with steps of 1 (for windows of one base) or 3. The most significant *p*-values are shown, found with the 1 base window. **D** Number of patients with each deleterious or significant missense mutations, distributed according to histology sub-type.

### Point mutations in EIF4G2 functional domains alter its protein interactome

To dissect the potential effects that the missense mutations have on EIF4G2 function, the six significant hot-spot missense somatic mutations were aligned to the domain organization scheme of the EIF4G2 protein (Fig 2A). While 35% of the EIF4G2 protein is predicted to consist of unstructured regions with unknown functions (e.g. predicted EIF4G2 structure, AlphaFold Protein Structure Database, https://alphafold.ebi.ac.uk/entry/P78344), it also contains 3 distinct functional domains whose crystal structures have been determined at high resolution (Fan *et al*, 2010; Liberman *et al*., 2008; Virgili *et al*, 2013). These include the MIF4G domain (aa positions 78-308), the mainly α-helical MA3-like domain (positions 543-666), and the C-terminal W2 domain (720-907). Two of the patient-derived mutations localized to the MIF4G domain (R178 and R295) (Fig 2A), which is known to interact with translation initiation factors EIF4A and EIF3, and also bind mRNA. As this implies a substantial contribution for this region to EIF4G2’s potential tumor functions, we also included in the analysis an additional missense mutation (R165) that was previously shown to be important for RNA binding (Virgili *et al*., 2013), despite the fact that it did not pass the significance score. The most abundant mutation position, R505, mapped to a large segment that is predicted to be unstructured with as-of-yet no known function. L545 is found at the start of the MA3-like domain, a region with unknown function. It is mostly buried (Fan *et al*., 2010) and mutation to F, with a larger side chain, may affect the local folding. The R714 mutation is located at the C-terminus of the MA3-like domain, just before the linker contacting it to the W2 domain (Fan *et al*., 2010). The N785 mutation mapped to the W2 domain that has been shown to interact with EIF5C, EIF2S2 and MNK1 (Asano *et al*, 1999; Liberman *et al*., 2008; Liberman *et al*., 2015; Pyronnet *et al*, 1999).

**Figure 2.**
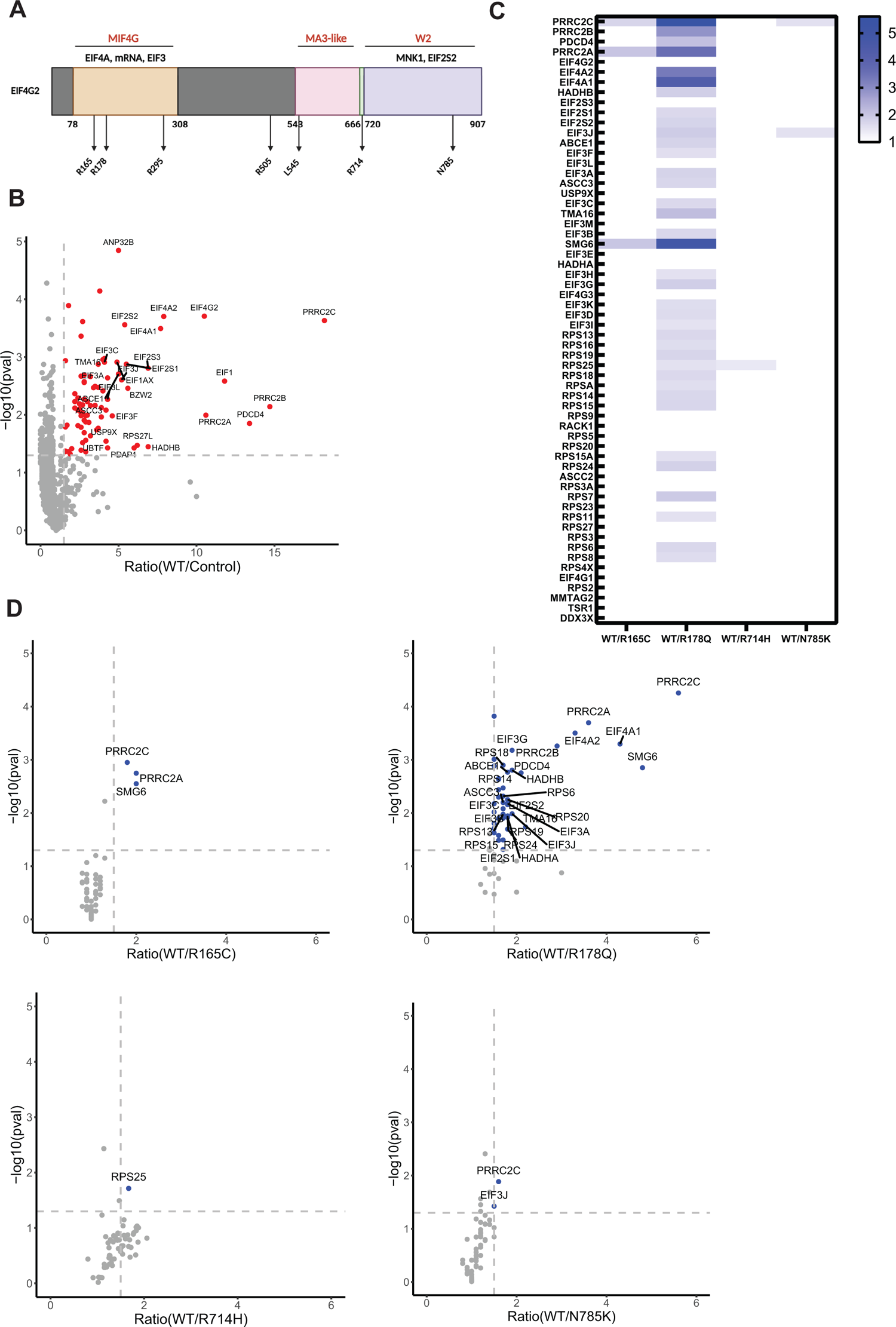
MIF4G point mutations reduce EIF4G2 protein-protein interactions. **A** Schematic representation of EIF4G2 protein. The domains with known crystal structure are designated by amino acid position and labeled in red. Proteins shown to interact with these regions are indicated in black. The approximate locations of the significant mutations are represented by arrows in the relevant domains. Scheme was created with BioRender.com. **B** Volcano plot of the fold-ratio of the abundance of the detected proteins in WT EIF4G2 vs control IP samples, vs. their significance expressed as -log_10_ *p*-value. Proteins with significant increased abundance, i.e., EIF4G2 interactors, are indicated in red. **C** Comparison of the binding abilities of EIF4G2 mutants to the 60 identified EIF4G2 interacting proteins. Heat map shows the fold-ratio of abundance of specific interacting proteins in the IP of the mutants compared to the WT EIF4G2 IP. The interactors presented are the ones identified as EIF4G2 interacting proteins (p<0.05, fold change>1.5 of WT/Control) regardless of their significance in the WT vs. mutant comparison, and are listed in order of their relative abundance in the WT vs control IP. Decreased interaction with the mutant EIF4G2 is represented by a stronger blue strip; white indicates no change in protein abundance. Only significant fold changes >1.5 with a *p*-value<0.05 are indicated, and only mutants for which such changes were observed are shown. **D** Volcano plot of the fold-ratio protein abundance in IPs of WT EIF4G2 compared to either R165C, 178Q, R714C, or N785K mutants, vs. their significance expressed as - log_10_ *p*-value. Only interacting proteins are shown. Proteins showing significant decreased abundance (>1.5 fold change, p<0.05) in the mutant IP relative to the WT IP are represented by blue dots.

Of these mutations, we chose to focus on the five-point mutations in the domains with established functions to investigate the mutations’ outcome on EIF4G2’s protein-protein interaction capability. The individual point mutations, chosen based on their abundance in the patient samples (i.e. R165C, R178Q, R295C, R714H, and N785K), were constructed in Flag-tagged EIF4G2, and transfected into HEK293T cells. Interactors were analyzed by co-immunoprecipitation [IP] followed by mass spectrometry analysis [MS] in two separate experiments, each comparing the protein interactome of WT EIF4G2 to either 3 or 2 of the mutants, respectively. EIF4G2 interactors were defined as those proteins that passed the threshold of a mean abundance greater than 1.5 compared to the control IP, with a *p*-value<0.05, in both MS runs. The first MS run yielded 83 interacting proteins (Fig 2B) while the second MS yielded 151 (Fig EV2A), with a shared set of 60 proteins; these 60 were defined as the set of EIF4G2 interactors. All but three have been previously linked to translation or mRNA regulation (Fig 2B, Fig EV2A, and Table EV1). 23 proteins were not previously reported by earlier studies of the protein interactome of EIF4G2 in mouse embryonic stem cells (Sugiyama *et al*, 2017) and MDA-MB-231 breast cancer cells (de la Parra *et al*., 2018a) (Fig EV2B). Yet consistent with these prior reports, the interactors mainly included components of the 43S pre-initiation complex [PIC], i.e., canonical translation initiation factors of the EIF3, EIF4A and EIF2S2 families and small ribosome subunits (20 and 24 proteins, respectively, Fig EV2C). In addition, the data confirmed the strong interaction with the PRRC2 proteins, recently implicated in translation initiation (Jiang *et al*, 2023). Several of the newly identified interacting proteins have been more indirectly linked to translation initiation, including ribosome assembly factors (i.e. TMA16 and TSR1 (Liang *et al*, 2020; Scaiola *et al*, 2018)), regulators of translation initiation (DDX3X (Ryan & Schroder, 2022), PDCD4 (Cai *et al*, 2022)) and several linked to mRNA surveillance and translation quality control, such as the endonuclease SMG6 that is involved in nonsense-mediated mRNA decay (Boehm *et al*, 2021), ABCE1 of the No-Go mRNA decay pathway (Nurenberg-Goloub & Tampe, 2019), and ASCC2, ASCC3 and RACK1, which mediate the ribosome-associated quality control pathway (Hashimoto *et al*, 2020; Joazeiro, 2017; Juszkiewicz *et al*, 2020) (Fig EV2C, Table EV1).

The R295C mutation in the MIF4G domain did not significantly alter the repertoire of EIF4G2 interacting proteins in the complex (Table EV1). In contrast, the R714H mutation near the W2 domain displayed a preferential significant decrease (i.e. >1.5-fold) in binding to only one interactor, RPS25, without affecting any of the other ribosomal proteins in the complex (Fig 2C,D). R165C, located in the MIF4G mRNA binding domain, showed a decreased interaction with three proteins, PRRC2A, PRRC2C, and SMG6 (Fig 2C, D), but did not impact on EIF4A or EIF3, which are known to bind to this domain. The C-terminal N785K mutant showed a significantly decreased interaction with PRRC2C and EIF3J (Fig 2C, D). The most dramatic effect on the abundance of the EIF4G2 interacting proteins in the co-IP was caused by the R178Q mutation. It significantly decreased the binding to 39 of the 60 EIF4G2 interacting proteins (Fig 2C, D), with the greatest impact on the interaction with all PRRC2 family members, followed by major loss of binding to the canonical translation initiation factors that are absolutely critical for EIF4G2 function in translation initiation, such as EIF4A, EIF3 and EIF2S members. At least part of this effect could be explained by the fact that the R178Q mutant was itself less abundant, albeit not significantly, in cell lysate compared to the WT protein (Table EV1, EV4). Nevertheless, all but one of the interacting proteins showed a fold-reduction in the mutant vs WT IPs that exceeded the fold-reduction of EIF4G2 itself (Table EV1). Moreover, the R714H mutant was even less abundant in the IP than the R178Q mutant, and yet did not show a significantly reduced interactome. Therefore, the changes in the binding of R178Q to its interactors are likely due to both changes in the protein steady-state levels and binding affinity. Thus, the patient-derived mutations in the known EIF4G2 functional domains have an impact on EIF4G2’s ability to interact with its protein partners in different ways. Furthermore, they hint to possible binding domains for those interactors that have not yet been identified and/or mapped to the EIF4G2 structure.

### MIF4G domain mutants show differential effects on IRES and uORF mediated mRNA translation

To determine whether the changes in protein binding observed for some of the mutants had an effect on EIF4G2 functional activity, and whether those with unchanged protein interaction profiles nevertheless lose functionality in other ways, cellular translation assays were performed. Various established EIF4G2 targets that represent different translation initiation mechanisms were assessed. These included BCL2, representing IRES-directed targets (Marash *et al*., 2008; Yoffe *et al*., 2016), and ROCK1 and WNK1, which contain cap-dependent uORFs that drive re-initiation of the downstream CDS in an EIF4G2-dependent mechanism (David *et al*., 2022; Weber *et al*., 2022). We first confirmed the EIF4G2-dependency of all three proteins in EIF4G2 knock-out (KO) 293T cells, generated by the CRISPR-Cas9 system (Fig EV3A). Western blotting indicated that deletion of EIF4G2 resulted in reduced protein steady state levels of endogenous BCL2, ROCK1 and WNK1. The three targets were then tested as reporters in a cellular dual luciferase translation system. The reporter for BCL2 consisted of an A-cap structure followed by a hairpin loop added upstream of the BCL2 IRES to ensure cap-independent initiation of a firefly luciferase [F-LUC] gene ((Liberman *et al*., 2015) and Fig 3A, scheme). For ROCK1 and WNK1 reporters, their entire 5’UTRs (including all uORFs) were inserted upstream of a Renilla luciferase [R-LUC] gene ((Weber *et al*., 2022), and schemes, Fig 3B and C). These were then co-transfected with either R-LUC or F-LUC, respectively, as internal controls, together with EIF4G2 WT or the panel of patient-derived mutants, in the EIF4G2 KO cells. The ability of the patient-derived mutants to drive translation of the reporters was compared to WT EIF4G2, after normalization to the second LUC control.

**Figure 3.**
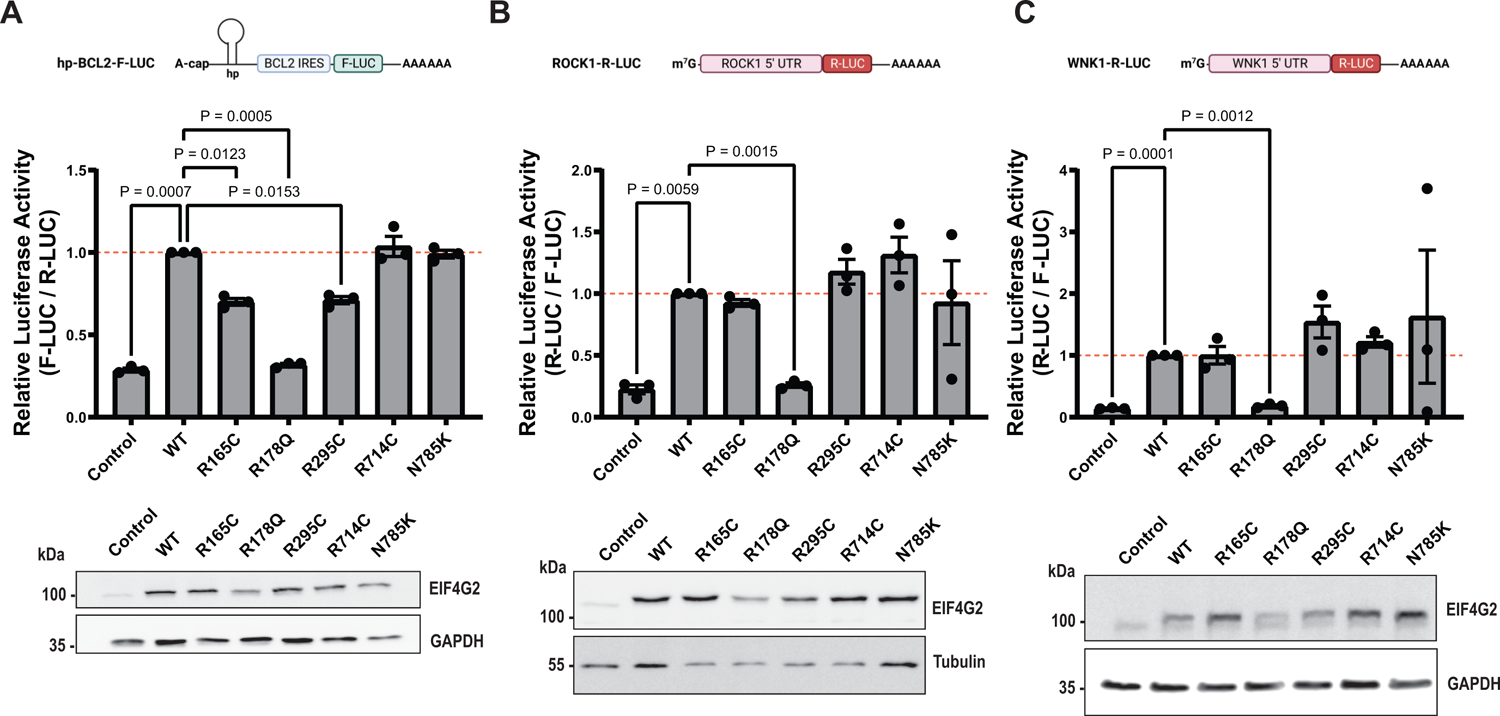
Effect of EIF4G2 mutations on IRES-dependent and uORF-dependent translation. **A** HEK293T EIF4G2 KO cells were co-transfected with the indicated EIF4G2 variants and an F-LUC reporter driven by the BCL2 IRES and R-LUC reporter as an internal control. A schematic of the F-LUC reporter is shown, including a mutant A-cap structure and a hairpin upstream of the IRES sequence. F-LUC activity was quantified and normalized to the R-LUC activity; the graph shows the relative normalized LUC activity in all EIF4G2 transfectants with WT EIF4G2 transfection set as 1 (dashed red line). Total cell lysates were subjected to western blot analysis using EIF4G2 and GAPDH antibodies as loading control, shown below the graph. **B,C** HEK293T EIF4G2KO cells were co-transfected with the indicated EIF4G2 variants and reporters containing ROCK1 5’UTR (**B**) or WNK1 5’UTR (**C**) upstream of R-LUC along with F-LUC as internal control. Schematics of the R-LUC reporters are shown. R-LUC activity was quantified and normalized to the F-LUC activity; the graph shows the relative normalized LUC activity in all EIF4G2 transfectants with WT EIF4G2 transfection set as 1 (dashed red line). Total cell lysates were subjected to western blot analysis using EIF4G2 and tubulin or GAPDH antibodies as loading controls, shown below the graphs. Data Information: For all panels, data is presented as individual data-points and also as the mean values±S.E.M of three (**A,C**) or four (**B**) independent experiments, with a representative western blot from one of the experiments shown. Significance was determined by matched one-way ANOVA followed by Dunnett’s multiple comparison ad hoc test (comparing all variants to the WT EIF4G2 construct). Non-significant results were not indicated in the figure. **A-C** Schemes were created with BioRender.com.

The R714C mutation and N785K mutation at the C-terminal domain had no effect on any of the luciferase reporters compared to the WT, indicating no functional significance for these translation initiation mechanisms (Fig 3A-C). We cannot, however, at this point, exclude the possibility that these mutants may be impaired in other functions not tested here, such as EIF3D-mediated cap-dependent initiation or m6A-driven translation. In contrast, the three MIF4G mutants all showed reduced activity towards the BCL2 IRES, but only one reduced the uORF-directed translation. Specifically, the R178Q mutant completely lost the ability to drive translation of the luciferase reporters for all targets tested; the LUC signals were reduced to the level of the control transfection (Fig 3A-C). The relative mRNA expression levels of the R-LUC and F-LUC reporters were assessed by quantitative PCR [qPCR] to exclude decreased reporter transfection or gene expression as causes for the observed decreased reporter activity (Fig EV3B, C). Thus, this mutant is incapable of driving either IRES-dependent or uORF-dependent initiation mechanisms. This is not surprising considering that the R178Q mutant showed reduced interactions with the canonical translation factors that are necessary for translation initiation. Furthermore, protein levels of the R178Q mutant were consistently lower than the WT and other mutants in all assays (Fig 3A-C), providing an additional explanation for its lost ability to drive translation of all tested targets.

In contrast, the R165C and R295C mutants significantly impaired the ability to translate the F-LUC reporter bearing the BCL2 IRES by 30% relative to the WT EIF4G2 (Fig 3A), without impacting on the translation of the ROCK1 and WNK1 reporters (Fig 3B, C). Notably, western blotting of lysates after co-transfection indicated that these mutants were expressed at similar or even greater levels compared to the WT EIF4G2, across the experiments, excluding any correlation between changes in protein expression and decreased activity (Fig 3A-C). Also, these decreases were not attributed to changes in mRNA expression of the reporters (Fig EV3C, D). Overall, the data imply a selective impact of these two mutations on IRES-mediated translation initiation and not translation directed by re-initiation from uORFs.

Of these two mutants that selectively reduced IRES-dependent translation, only the R165C mutation had any effect on EIF4G2 protein interactions, reducing its ability to bind PRRC2A, PRRC2C and SMG6. In contrast, the interaction of the R295C mutant with the entire cohort of EIF4G2 interacting proteins was not changed. Thus, it is unlikely that the functional effects that these mutations have on IRES-dependent translation is due to changes in protein binding. As the MIF4G domain has been previously shown to mediate RNA binding, including, specifically, the R165 position, it is more likely that these mutations specifically affect binding to target mRNAs bearing IRESes. In fact, EIF4G2 has been shown to be capable of directly binding mRNA targets with IRESes, such as p53 and HMGN3, in electophoretic mobility shift assays (Weingarten-Gabbay *et al*., 2014; Yoffe *et al*., 2016). It is reasonable to assume that binding to IRES-dependent targets differs in some inherent manner from binding to other targets, such as those that contain uORFs. The fact that 2 specific point mutations within the mRNA binding domain affect translation of an EIF4G2 IRES target but not uORF-dependent targets indicates that the protein-RNA interaction for the former is more sensitive to these subtle changes in the EIF4G2 structure, and serves as the first clue to potentially explain preferential target recognition.

### 3D structure analysis predicts loss of protein interactions by MIF4G and W2 domain mutants

In order to understand why the mutations impact on EIF4G2’s ability to bind its interacting proteins and promote translation, 3D structural analysis was performed. Model structures of the complete EIF4G2 generated by AlphaFold (https://alphafold.ebi.ac.uk/entry/P78344) or by roseTTAfold (Baek *et al*, 2021) gave conflicting predictions as to the relative organization of the known domains and unstructured regions. We therefore focused on the experimentally solved structures. The structure of the W2 domain (Fan *et al*., 2010) shows that residues R714 and N785 lie on the surface, and thus mutations of these positions to His and Lys, respectively, may affect intermolecular interactions (Fig 4A). A deep cavity is located near N785, lined with hydrophobic aromatic and aliphatic residues, which may be a protein-protein binding pocket.

**Figure 4.**
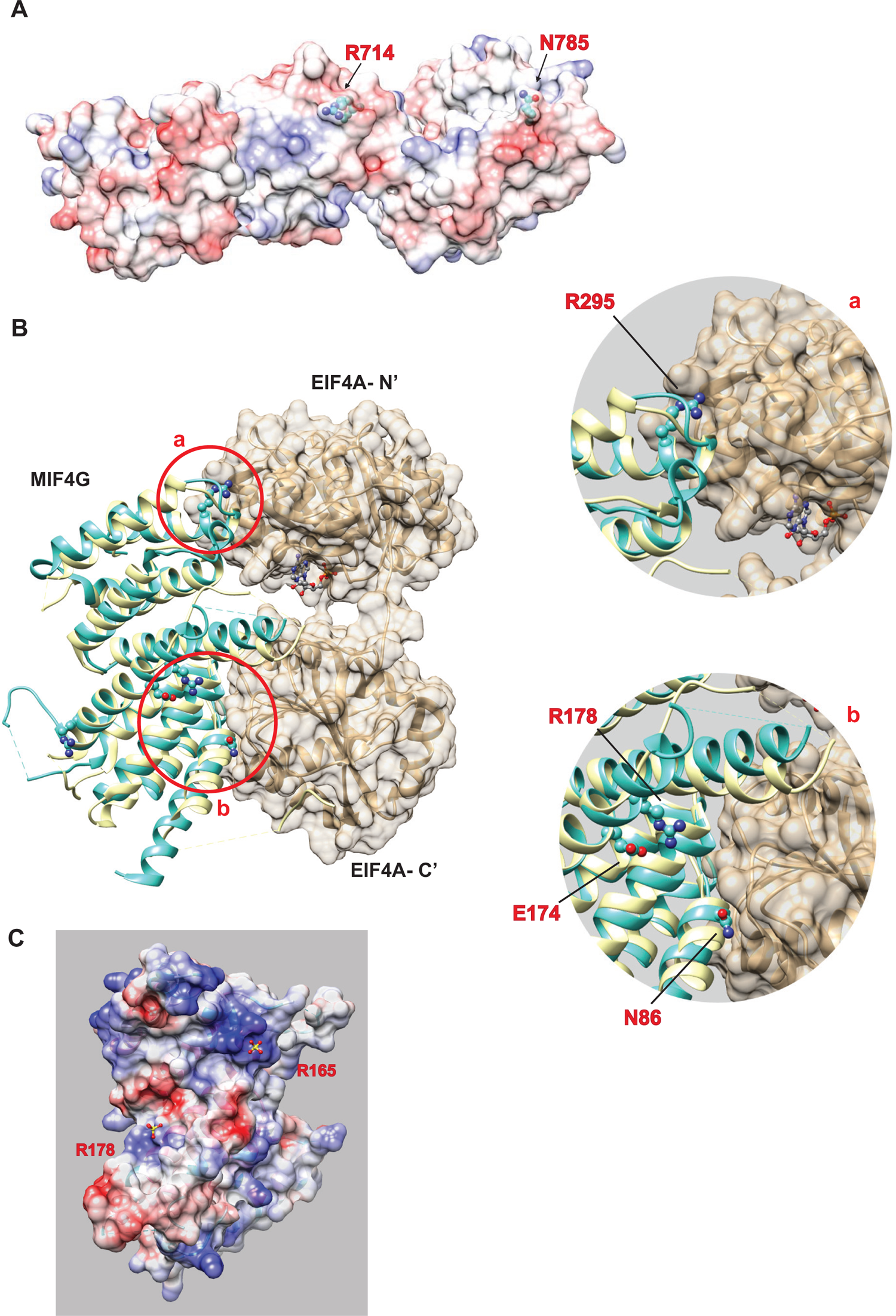
EIF4G2 structure analysis predicts possible outcome of patient-derived significant missense mutations. **A** The surface of the C-terminal domains of EIF4G2 (PDB entry 3L6A) showing the positions of mutations R714H and N785T/K. The surface is colored by the Coulombic potential, red for negative, blue for positive and white for neutral. **B** Model of the interaction between the MIF4G domain of EIF4G2 and EIF4A based on the structure of the yeast complex between EIF4G (yellow) and EIF4A (beige) (PDB entry 2VSX). The structure of human EIF4G2 MIF4G domain (cyan) (PDB entry 4IUL) was superposed on yeast EIF4G. As in the yeast complex, MIF4G interacts with two domains of EIF4A. **Upper inset,** magnification of the circled area a, showing the interface with the N-terminal domain of EIF4A. R295 (K837 in yeast EIF4G) is marked in red. **Lower inset**, magnification of the circled area b, showing the interface with the C-terminal domain of EIF4A, highlighting the position of N86 (N86 corresponds to N615 in yeast eIF4G). R178 (K709 in yeast) is shown, making an ion pair with E174 (E706 in yeast). **C** The surface of DAP5 MIF4G domain (PDB entry 2VSX), showing a trough between R165 and R178. The surface is colored by the Coulombic potential, blue for positive, red for negative and white for neutral. Sulfate ions are shown as ball-and-stick, yellow and red, respectively, for the sulfur and oxygen atoms. The sulfate ion near R165 is located within a deep and strongly positive cavity. The sulfate ion near R178 is located near a weakly positive surface region.

For the structure analysis of the EIF4G2 MIF4G domain and its interacting partner EIF4A, the structure of human EIF4G2 MIF4G domain ((Virgili *et al*., 2013) was superimposed on the yeast EIF4G MIF4G-EIF4A ((Schutz *et al*, 2008) (Fig 4B). In the crystal structure, residue EIF4G K837, corresponding to human EIF4G2 R295, interacts electrostatically with the N-terminal domain of EIF4A (Fig 2B, upper inset). Residue EIF4G N615, corresponding to human EIF4G2 N86, interacts with the C-terminal domain of EIF4A (Fig 2B, lower inset); N86 is a well-studied position that is critical for EIF4A binding (Liberman *et al*., 2015; Virgili *et al*., 2013). Both interactions are necessary to control the relative orientation of the two domains of EIF4A. The R295C mutation likely weakens the electrostatic interaction with EIF4A and possibly affects the relative position of the two domains of EIF4A, and consequently, mRNA binding and/or translation. R165 is located near a deep cavity with strong positive electrostatic charge, and serves as an mRNA binding site together with K108 and K112 (Virgili *et al*., 2013) (Fig 4C). The R165C mutation may neutralize the positive electrostatic surface, and subsequently, weaken the affinity for target mRNAs that rely on this region for binding, ultimately reducing the efficiency of their translation. The R178Q mutation had the most dramatic effects on the EIF4G2 protein, reducing both its ability to bind its interacting partners and to drive translation of all tested targets. These global loss-of-function effects most likely are related to the fact that its steady state levels when over-expressed were vastly reduced compared to WT EIF4G2 (Fig EV4). As for possible effect on interactions, we note that, although the positive charge of R178 is partially offset by the ion pairing with E174, the surrounding surface is moderately positive (Figure 4C). The mutation R178C eliminates the positive side chain and precludes the ion pairing, resulting in neutralized surface potential. The structural analysis of the MIF4G domain alone did not provide an explanation for why the mutation would cause a major reduction in the steady-state levels or loss of functionality, but in the absence of an experimental structure of the entire protein, intra-molecular contacts and overall protein folding could not be assessed.

In conclusion, we have identified several EIF4G2 somatic mutations in primary tumors of cancer patients that show various impairments in binding to interacting proteins and ability to direct mRNA translation. This correlation between impaired protein function and cancer is promising in establishing EIF4G2 as a potential tumor suppressor, at least in early pre-metastatic stages. Follow-up experiments specifically examining potential tumor-promoting attributes of these variants in cells and/or animal models are mandated to determine whether in fact loss of EIF4G2 function contributes to tumor development or growth.

## Materials and Methods

### Data collection and mutation analysis of EIF4G2 gene in human cancer

Whole genome screens data from the Catalogue of Somatic Mutations in Cancer, COSMIC (http://cancer.sanger.ac.uk/cancergenome/projects/cosmic), version 96-38, was analyzed for somatic mutations of the EIF4G2 gene in human cancer as previously described (Nuta *et al*., 2019). In brief, unique independent samples were identified by comparing all their listed mutations. Confirmed somatic mutations of the EIF4G2 gene were classified by mutation type. Further analysis of the coding regions mutations calculated the significance of each cluster of mutations by statistics of observed vs. expected mutations using a Poisson distribution for different nucleotide intervals.

### TCGA data mining

Mining of EIF4G2 mRNA expression levels in healthy and primary tumor samples was done using the UCSC Xena Functional Genomic Explorer (https://doi.org/10.1038/s41587-020-0546-8). All the chosen studies were TCGA based, followed by search of genomic EIF4G2 gene expression data, focusing on the sample_type phenotype.

### Cell Lines and Cell Culture

HEK293T cells (ATCC CRL-3216) was cultured in DMEM medium (Biological Industries) with 10% FBS (Gibco), 1% penicillin streptomycin (Biological Industries) and 1% L-Glutamine (Biological Industries). Cells were routinely screened for mycoplasma.

### Structural analysis

Molecular graphics and structure analyses were performed with UCSF Chimera, developed by the Resource for Biocomputing, Visualization, and Informatics at the University of California, San Francisco, with support from NIH P41-GM103311 (Pettersen *et al*, 2004).

### Generation of EIF4G2 KO cells and EIF4G2 mutant plasmids

HEK293T EIF4G2 KO cells were generated by the CRISPR-Cas9 method, targeting exon 7 of the CDS. The following guides were cloned into pKLV vector (kindly gifted by Y. Shaul, Weizmann Institute of Science, Israel): EIF4G2 sense: CACCATTAGACCATGAACGAGCC, EIF4G2 antisense: TAAAACTGGCTCGTTCAGGTCT. 293T cells were transfected by standard calcium-phosphate transfection reagents. After 48 h, the transfected cells were treated with puromycin (Sigma-Aldrich) for selection of the transfected cells. The puromycin-resistant cells were transferred by limited dilution into a 96-well plate, then expanded and screened for the presence of the CRISPR KO by colony PCR followed by DNA sequencing and western blot for validation.

EIF4G2 mutants were created from the WT Flag-tagged EIF4G2 template in pcDNA3 (pcDNA3-FLAG-EIF4G2_WT (Liberman *et al*., 2015)), using standard transfer-PCR or point mutagenesis cloning procedures with the primers listed in Table 1.

**Table 1.**
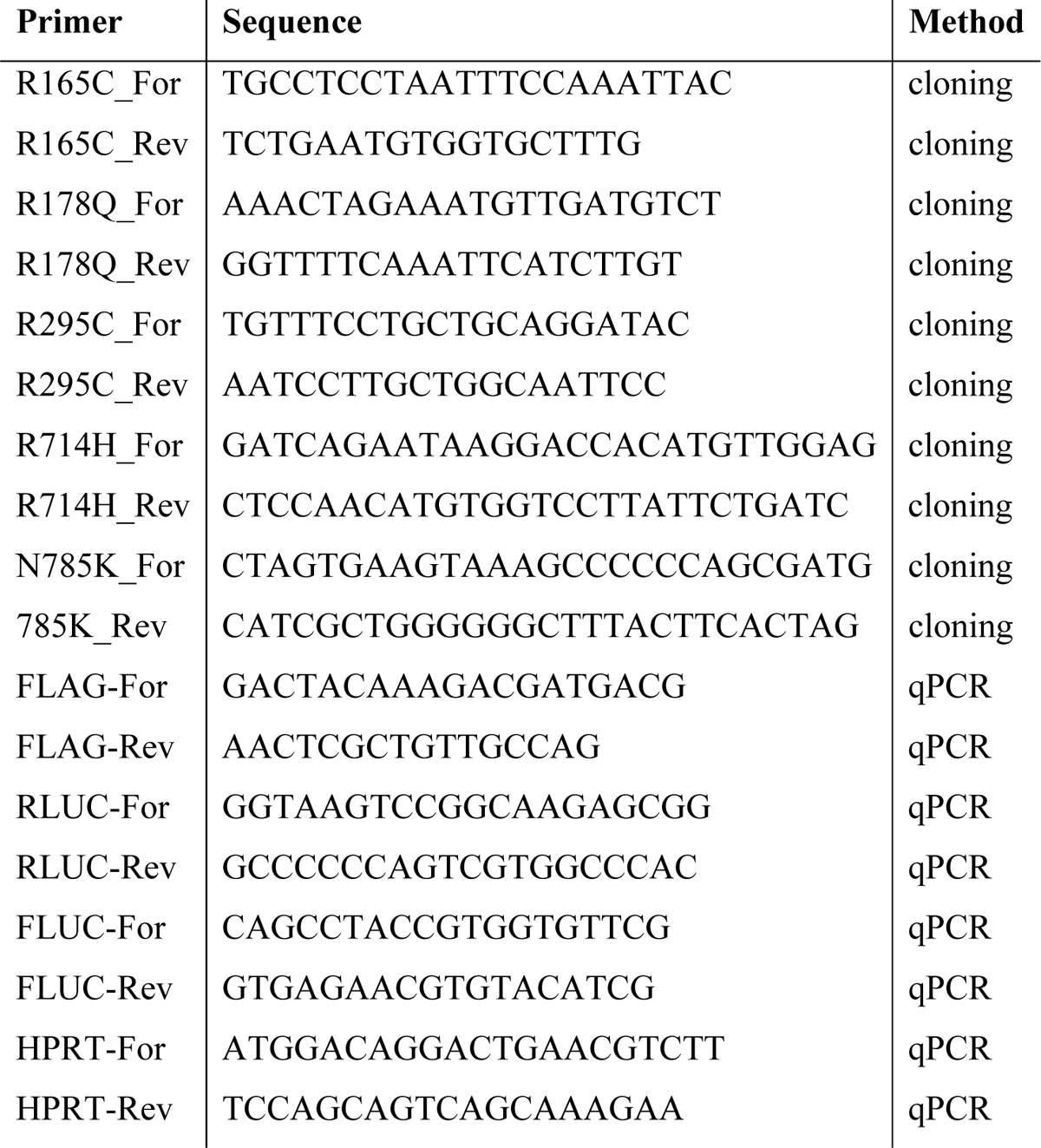
List of primers use for PCR cloning and qRT-PCR.

### Co-immunoprecipitation and Mass Spectrometry

HEK293T cells were transfected with the following plasmids in two separate biological experiments by standard calcium-phosphate transfection: the first consisted of pcDNA3-FLAG-mCherry, pcDNA3-FLAG-EIF4G2_WT, pcDNA3-FLAG-EIF4G2_R165C, pcDNA3-FLAG-EIF4G2_R178Q and pcDNA3-FLAG-EIF4G2_R295C, and the second consisted of the FLAG-mCherry and EIF4G2 WT plasmids, and also pcDNA3-FLAG-EIF4G2_R714H, and pcDNA3-FLAG-EIF4G2_N785K. After 48 h, the cells were harvested and lysed in B-buffer (20 mM HEPES-KOH [pH 7.6], 100 mM KCl, 0.5 mM EDTA, 0.4% NP-40, 20% glycerol) supplemented with protease and phosphatase inhibitors (Sigma-Aldrich). 5 mg lysates were incubated with FLAG beads (Sigma-Aldrich, cat# A2220) at 4°C for 2 h; a small amount was reserved for western blot validation of protein expression. The beads were washed 3 times with lysis buffer and the bound protein was eluted using SDS buffer (5% SDS in 100mM Tris-HCl, pH 7.4).

20 μg protein from the eluted samples of each experiment were subjected separately to in-solution tryptic digestion using the suspension trapping (S-trap micro-columns, Protifi) method as previously described (Elinger *et al*, 2019). For the first IP experiment (Control, WT EIF4G2, R165C, R178Q and R295C mutants), the resulting peptides were loaded using split-less nano-Ultra Performance Liquid Chromatography (10 kpsi nanoAcquity; Waters, Milford, MA, USA). The peptides were separated using an Aurora column (75μm ID x 25cm, IonOpticks) at 0.3 µL/min. Peptides were eluted from the column into the mass spectrometer using the following gradient: 2% to 30%B in 41 min, 30% to 90%B in 2 min, maintained at 90% for 3 min and then back to initial conditions. The nanoUPLC was coupled online to a quadrupole orbitrap mass spectrometer (timsTOF Pro, Bruker). Data was acquired in data dependent acquisition with ion mobility mode (DDA PASEF (Meier *et al*, 2018)), using a 1.1sec cycle-time method with 10 MS/MS scans. For ion mobility 1/K0 range was 0.60-1.60 Vs/cm^2^, Energy Start in PASEF CID was set to 20.0 ev and Energy End was set to 59.0 eV. Other parameters were kept as the default parameters of the DDA PASEF method. The raw data was processed with FragPipe v17.1. The data was searched with the MSFragger search engine v3.4 against the human (Homo sapiens) protein database as downloaded from Uniprot.org, appended with common lab protein contaminants. Enzyme specificity was set to trypsin and up to two missed cleavages were allowed. Fixed modification was set to carbamidomethylation of cysteines and variable modification was set to oxidation of methionines, and protein N-terminal acetylation. The quantitative comparisons were calculated using Perseus v1.6.0.7. Decoy hits were filtered out and only proteins that had at least 2 valid values after logarithmic transformation in at least one experimental group were kept. For statistical calculations missing values were replaced by random values from a normal distribution using the Imputation option in Perseus (width 0.3, downshit 1.8). A Student’s t-test of the logarithmic transformation was used to identify significant differences between the experimental groups, across the biological replica. Fold changes were calculated based on the ratio of geometric means of the different experimental groups.

For the second IP, experiment (Control, WT EIF4G2, R714H and N785K mutants), each sample was loaded using split-less nano-Ultra Performance Liquid Chromatography as above, except that desalting of the samples was performed online using a reversed-phase Symmetry C18 trapping column (Waters) and the peptides were then separated using a T3 HSS nano-column (Waters) at 0.35 µL/min. Peptides were eluted from the column into the mass spectrometer using the following gradient: 4% to 27%B in 55 min, 27% to 90%B in 5 min, maintained at 90% for 5 min and then back to initial conditions. The nanoUPLC was coupled online through a nanoESI emitter (10 μm tip; New Objective; Woburn, MA, USA) to a quadrupole orbitrap mass spectrometer (Q Exactive HFX, Thermo Scientific) using a FlexIon nanospray apparatus (Proxeon). Data was acquired in data dependent acquisition (DDA) mode, using a Top10 method. MS1 resolution was set to 120,000 (at 200m/z), mass range of 375-1650m/z, AGC of 1e6 and maximum injection time was set to 50 ms. MS2 resolution was set to 15,000, quadrupole isolation 1.7m/z, AGC of 1e5, dynamic exclusion of 20 s and maximum injection time of 60 ms. Raw data was processed with the MetaMorpheus algorithm version 0.0.320 (Solntsev *et al*, 2018). The data was searched against the human (Homo sapiens) protein database as downloaded from Uniprot (www.uniprot.com), and appended with common lab protein contaminants. Enzyme specificity was set to trypsin and up to two missed cleavages were allowed. Fixed modification was set to carbamidomethylation of cysteines and variable modification was set to oxidation of methionines. Peptide and protein identifications were filtered at an FDR of 1%. The minimal peptide length was 7 amino-acids. Peptide identifications were propagated across samples using the match-between-runs option checked. Searches were performed with the label-free quantification option selected. The quantitative comparison and statistics were calculated as above.

EIF4G2 binding proteins were defined as those with abundance at least 1.5 orders of magnitude greater in the WT EIF4G2 IP compared with the control Flag-Cherry IP sample, with *p*<0.05, common to both IP-MS experiments. For these proteins, the relative abundance compared to the WT EIF4G2 IP was calculated for each EIF4G2 mutant IP; loss of binding was defined as at least a 1.5-fold decrease in the fold ratio of mutant vs WT EIF4G2 IPs, with *p*<0.05.

### Western Blot

Cells were lysed in RIPA lysis buffer (20 mM Tris, pH 8.5, 0.1% NP40, 150mM NaCl, 0.5% sodium deoxycholate, 0.1% SDS) supplemented with 10 µl/ml 0.1M PMSF (Sigma-Aldrich, 93482) and 1% protease inhibitor (Sigma-Aldrich, P8340), or alternatively, directly in SDS sample buffer (300 mM Tris-Cl pH 6.8, 0.6% glycerol, bromophenol blue, 600 mM β-mercaptoethanol, 400 μM SDS). The former buffer was supplemented with 10 µl/ml 0.1M PMSF and 1% protease inhibitor (Sigma-Aldrich). Proteins were separated by SDS-PAGE and transferred to nitrocellulose membranes, which were incubated with the indicated antibodies: mouse anti-EIF4G2 (BD Biosciences, cat# 610742, RRID:AB_398065, 1:1000 dilution), mouse anti-FLAG (Sigma-Aldrich, cat# F3165, RRID:AB_259529, 1:1000), mouse anti-GAPDH (Millipore, cat# MAB374, RRID:AB_2107445, 1:3000), rabbit anti-ROCK1 (Cell Signaling Technology, cat# 4035, RRID:AB_2238679, 1:500), rabbit anti-WNK1 (Cell Signaling Technology, cat# 4979, RRID:AB_2216752, 1:500) and mouse anti-Tubulin (Sigma-Aldrich, cat# T9026, RRID:AB_477593, 1:70000) add BCL2 (Santa Cruz Biotechnology Cat# sc-509, RRID:AB_626733, 1:1000). Secondary antibodies consisted of either horseradish peroxidase (HRP)-conjugated goat anti-mouse (Jackson ImmunoResearch Labs, cat# 115-035-003, RRID: AB_10015289) or anti-rabbit (Jackson ImmunoResearch, cat# 111-165-144), which were detected by enhanced chemiluminescence using EZ-ECL (Biological Industries).

### Luciferase Translation Assay

2×10^5^ 293T-EIF4G2KO cells were seeded in 6 well plates and transfected with 5 μg empty pcDNA3, pcDNA3-FLAG-EIF4G2_WT, pcDNA3-FLAG-EIF4G2_R165C, pcDNA3-FLAG-EIF4G2_R178Q, pcDNA3-FLAG-EIF4G2_R295C, pcDNA3-FLAG-EIF4G2_R714H or pcDNA3-FLAG-EIF4G2_N785K together with 1 μg pHP-IRES-BCL2-F-LUC plasmid (Liberman *et al*., 2015; Yoffe *et al*., 2016) with 1 μg pCIneo-R-LUC plasmid as control, or alternatively, 1 μg pCIneo-WNK1-R-LUC or pCIneo-ROCK1-R-LUC with 1μg pEGFP-N3-F-LUC plasmids (kindly gifted from the Igreja lab (Weber *et al*., 2022), Max Planck Institute, Germany), using Lipofectamine 2000 (Invitrogen) according to the manufacturer’s recommendations.

48 h following transfection, cells were washed and harvested. Cell pellet was divided for luciferase activity, western blot and qPCR analysis. For dual luciferase assay, cells were lysed and luciferase activity was measured on identical quantities of lysate according to manufacturer’s guidelines, using substrates for both F-LUC and R-LUC in sequential reactions. Luciferase signal was read using a Veritas microplate luminometer (Turner Biosystems). R-LUC/F-LUC or F-LUC/R-LUC activity of the control and mutants were normalized to that of WT EIF4G2.

### Reverse transcription (RT) and quantitative PCR (qPCR)

0.5 μg RNA was mixed with 4 μL 5X reaction buffer and 1 μL RTase (AzuraQuant cDNA synthesis kit, Azura Genomics cat: AZ1996). Reaction mix was incubated at 42°C for 30 min and denatured at 85°C for 10 min. The reaction was stopped by incubating the samples at 10°C for 10 min. Real time qPCR (qPCR) was performed using 166.6 ng cDNA, 10μM forward and reverse primers (see Table 1) and 5μL of AzuraView GreenFast qPCR Blue Mix LR (Azura Genomics, AZ2305). Data was analyzed using QuantStudio 5 software (Thermo-Scientific). mRNA levels were normalized to housekeeping gene (HPRT) and the quantifications were calculated using the Livak method.

### Data Availability

The mass spectrometry proteomics data have been deposited in the MassIve repository of the ProteomeXchange consortium (https://massive.ucsd.edu), with the dataset identifier MSV000092704.

### Statistical Analysis

Statistical analysis was done using GraphPad Prism 9. Data is presented as the mean values±S.E.M of independent experiments. One-way matched ANOVA with Dunnett’s multiple comparison post-hoc tests or paired two-tailed t-tests were performed as indicated in the figure legends, with p<0.05 considered statistically significant.

## Acknowledgements

We would like to thank Nadav Goldberg for his advice and many fruitful discussions. This research was supported by a grant from the Pasteur-Weizmann Council and from the Institut Pasteur and the Weizmann Institute of Science.

## Conflict of Interest

The authors have no competing interests to declare.

## Author Contributions

**S.M.** conceptualized experimental design, performed experiments, interpreted results, analyzed data, wrote the manuscript, and prepared figures. **M.B.** performed experiments, interpreted results. **M.E** modeled and analyzed 3D protein structure. **A.S., Y.L.** performed the MS data acquisition and analysis. **S.B.** interpreted results, analyzed data, wrote the manuscript, prepared figures. **S.P.** performed and analyzed mutations in cancer databases. **A.K.** conceptualized experimental design, supervised the project, analyzed data, edited the manuscript, and provided funding. All authors read, edited and approved the final submitted manuscript.

## Expanded View Figure Legends

**Figure EV1** *related to Fig 1*

**A** EIF4G2 Log2 fold-change of normalized read count from healthy and primary tumor samples, as sequenced in 24 cancer histology sub-types. Data was collected from the UCSC Xena Functional Genomic Explorer. Significance was assessed using two tailed Student’s t-test (*, *p*<0.05; **, *p*<0.01; ***, *p*<0.001).

**B** Table listing all the histology sub-type names as presented in Figure EV1A and in the paper.

**Figure EV2** *related to Fig 2*

**A** Volcano plot of the second MS experiment, of the fold-ratio of the abundance of the detected proteins in WT EIF4G2 vs control IP samples, vs. their significance expressed as -log_10_ *p*-value. Proteins with significant increased abundance, i.e., EIF4G2 interactors, are indicated in red.

**B** Venn diagram showing overlap with previously reported DAP5 IP-MS data. 118 significant interactors from IPs of EIF4G2 from mESCs ((Sugiyama *et al*., 2017), Table S1, all proteins enriched over control IP with *p*<0.05) and 82 significant interactors from MDA-MB-231 breast cancer cells ((de la Parra *et al*., 2018a), Supplementary Data 2, all proteins enriched over control IP with FDR<0.05).

**C** Pie chart showing gene annotation of protein interactors of EIF4G2 identified in current study in HEK293T cells.

**Figure EV3** *related to Fig 3*

**A** Total cell lysate from HEK293T WT and EIF4G KO cells were subjected to western blot analysis using EIF4G2 and the indicated antibodies to EIF4G2 targets. Each protein was run separately and its respective loading control, Tubulin or GAPDH, is shown below each respective blot. Shown are representative blots of 3 independent experiments.

**B, C** Quantitative PCR (qPCR) analysis was performed on mRNA extracted from the same lysates used in Fig 3B and C, respectively. Shown is the relative normalized LUC expression (R-LUC/F-LUC) in all samples, with WT EIF4G2 transfection set as 1 (dashed red line).

Data Information: For (B, C) data is presented as individual data points and as the mean values±SEM of three independent experiments. Significance was determined by matched one-way ANOVA test followed by Dunnett’s multiple comparison ad hoc test, comparing all samples to WT.

**Figure EV4** *related to Fig 4*

Western blot of EIF4G2 KO HEK293T cells expressing Flag tagged WT or R178Q EIF4G2. Tubulin was used as a loading control. EIF4G2 signal was normalized to Tubulin and quantification results are represented as individual data points and also as mean values±SEM of 3 independent experiments. Significance was determined by two tailed t-test.

## Notes

### Competing Interest Statement

The authors have declared no competing interest.

### Summary of Updates

Addition of supplementary figures

