## Supplementary Figures for "Loss-of-function cancer-associated mutations in the EIF4G2 non-canonical translation initiation factor"

### Meril, et al. Supplementary Information- Figures

#### Meril et al. Expanded View 1

A

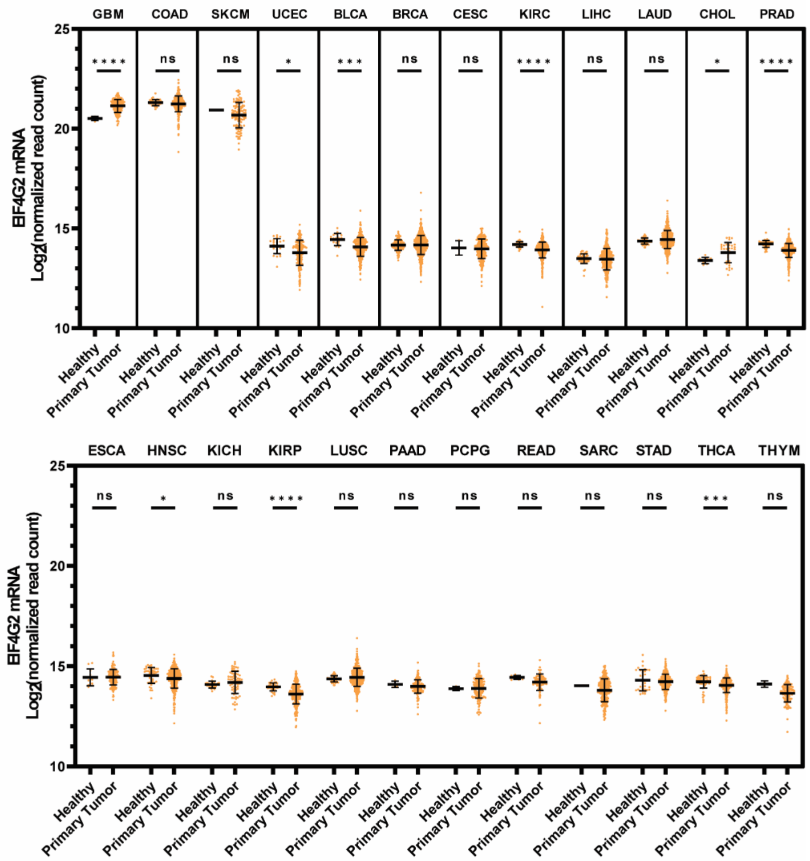


B

| TCGA Short name | Detailed Name |
|---|---|
| BLCA | Bladder Urothelial Carcinoma |
| BRCA | Breast invasive carcinoma |
| CESC | Cervical squamous cell carcinoma and endocervical adenocarcinoma |
| CHOL | Cholangio carcinoma |
| COAD | Colon adenocarcinoma |
| ESCA | Esophageal carcinoma |
| GBM | Glioblastoma multiforme |
| HNSC | Head and Neck squamous cell carcinoma |
| KICH | Kidney Chromophobe |
| KIRC | Kidney renal clear cell carcinoma |
| KIRP | Kidney renal papillary cell carcinoma |
| LIHC | Liver hepatocellular carcinoma |
| LUAD | Lung adenocarcinoma |
| LUSC | Lung squamous cell carcinoma |
| PAAD | Pancreatic adenocarcinoma |
| PCPG | Pheochromocytoma and Paraganglioma |
| PRAD | Prostate adenocarcinoma |
| READ | Rectum adenocarcinoma |
| SARC | Sarcoma |
| SKCM | Skin Cutaneous Melanoma |
| STAD | Stomach adenocarcinoma |
| THCA | Thyroid carcinoma |
| THYM | Thymoma |
| UCEC | Uterine Corpus Endometrial Carcinoma |

**Figure EV1** *related to Fig 1*

**A** EIF4G2 Log2 fold-change of normalized read count from healthy and primary tumor samples, as sequenced in 24 cancer histology sub-types. Data was collected from the UCSC Xena Functional Genomic Explorer. Significance was assessed using two tailed Student's t- test (*, $p<0.05$; **, $p<0.01$; ***, $p<0.001$).

**B** Table listing all the histology sub-type names as presented in Figure EV1A and in the paper.

**Meril et al. Expanded View 2**

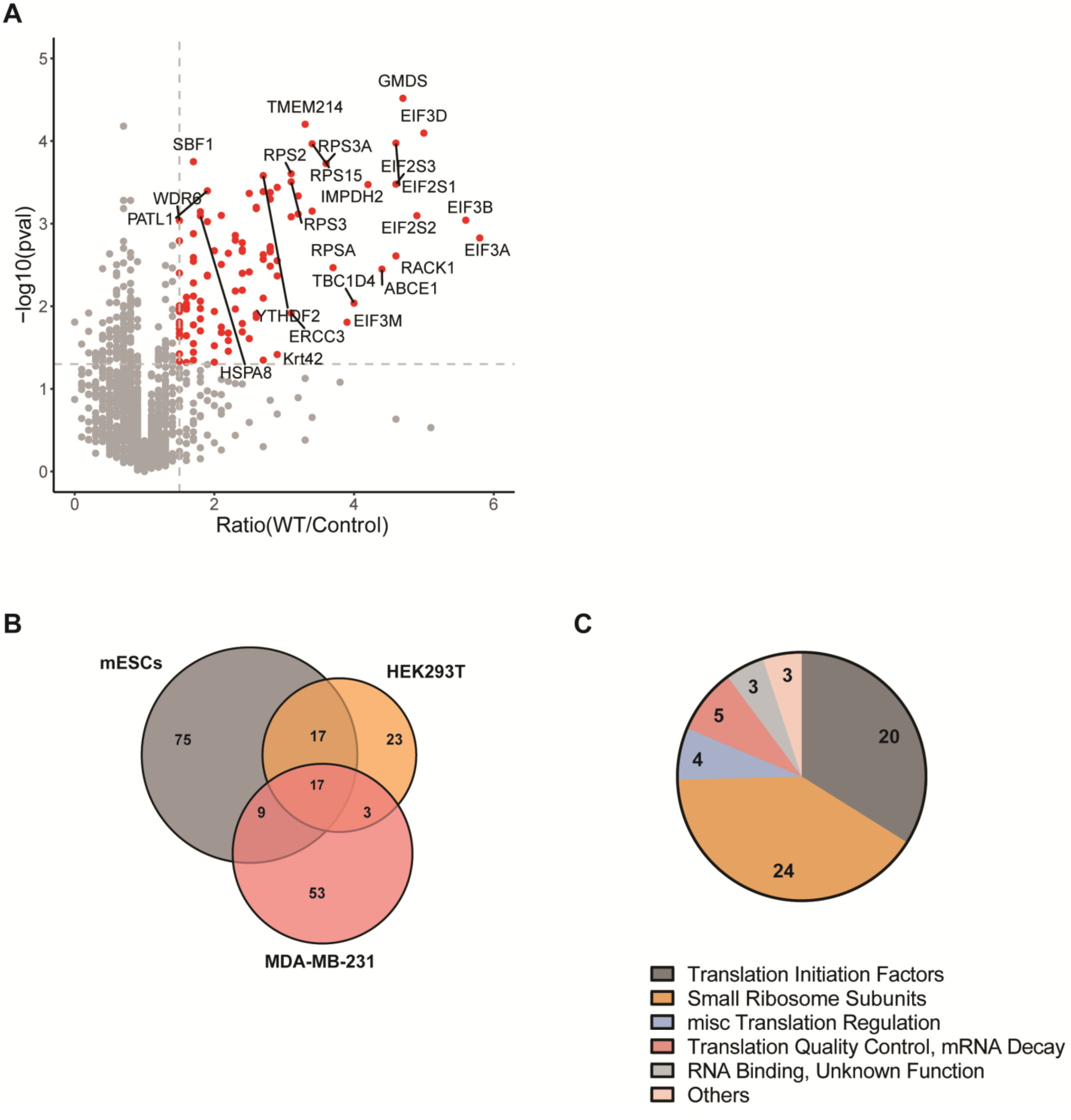

**Figure EV2** *related to Fig 2*

**A** Volcano plot of the second MS experiment, of the fold-ratio of the abundance of the detected proteins in WT EIF4G2 vs control IP samples, vs. their significance expressed as $-\log_{10}$ *p*-value. Proteins with significant increased abundance, i.e., EIF4G2 interactors, are indicated in red.

**B** Venn diagram showing overlap with previously reported DAP5 IP-MS data. 118 significant interactors from IPs of EIF4G2 from mESCs ((Sugiyama *et al.*, 2017), Table S1, all proteins enriched over control IP with $p<0.05$) and 82 significant interactors from MDA-MB-231 breast cancer cells ((de la Parra *et al.*, 2018a), Supplementary Data 2, all proteins enriched over control IP with FDR<0.05).

**C** Pie chart showing gene annotation of protein interactors of EIF4G2 identified in current study in HEK293T cells.

#### Meril et al. Expanded View 3

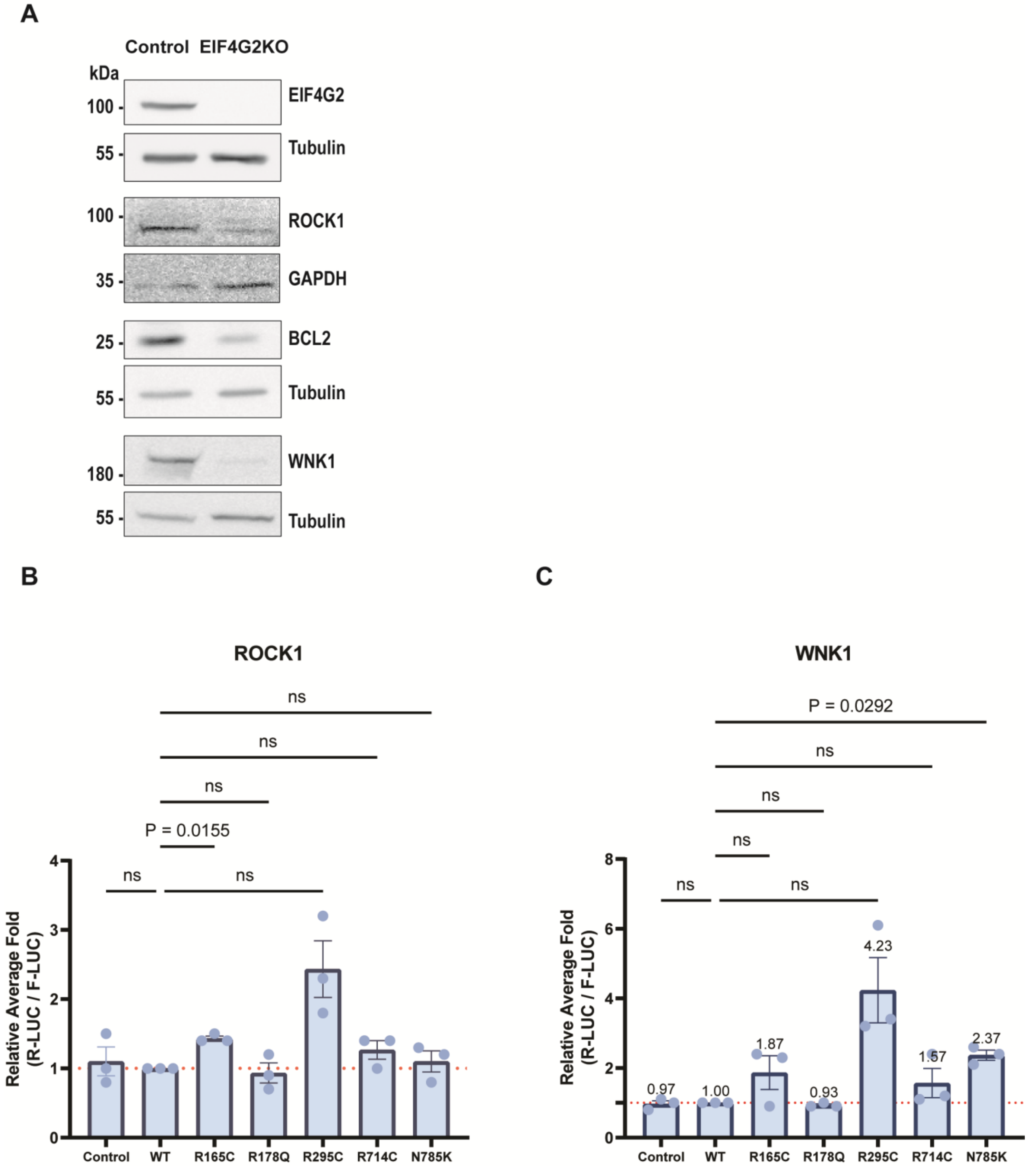

**Figure EV3** *related to Fig 3*

**A** Total cell lysate from HEK293T WT and EIF4G KO cells were subjected to western blot analysis using EIF4G2 and the indicated antibodies to EIF4G2 targets. Each protein was run separately and its respective loading control, Tubulin or GAPDH, is shown below each respective blot. Shown are representative blots of 3 independent experiments.

**B, C** Quantitative PCR (qPCR) analysis was performed on mRNA extracted from the same lysates used in Fig 3B and C, respectively. Shown is the relative normalized LUC expression (R-LUC/F-LUC) in all samples, with WT EIF4G2 transfection set as 1 (dashed red line).

Data Information: For (B, C) data is presented as individual data points and as the mean values±SEM of three independent experiments. Significance was determined by matched one-way ANOVA test followed by Dunnett’s multiple comparison ad hoc test, comparing all samples to WT.

### Meril et al. Expanded View 4

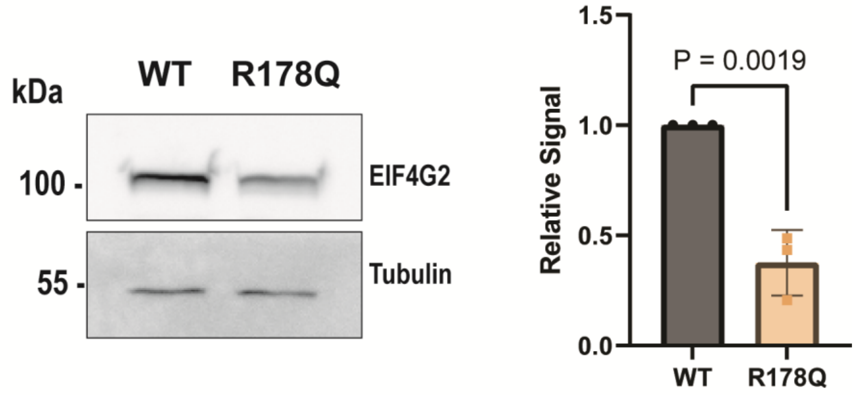


**Figure EV4** *related to Fig 4*

Western blot of EIF4G2 KO HEK293T cells expressing Flag tagged WT or R178Q EIF4G2. Tubulin was used as a loading control. EIF4G2 signal was normalized to Tubulin and quantification results are represented as individual data points and also as mean values±SEM of 3 independent experiments. Significance was determined by two tailed t-test.
